# The complete mitochondrial genome of the Golden Birdwing butterfly *Troides aeacus*: insights into phylogeny and divergence times of the superfamily Papilionoidea

**DOI:** 10.1101/529347

**Authors:** Si-Yu Dong, Guo-Fang Jiang, Gang Liu, Fang Hong, Yu-Feng Hsu

**Affiliations:** College of Oceanology & Food Science, Quanzhou Normal University, Quanzhou, China; Jiangsu Key Laboratory for Biodiversity and Biotechnology, College of Life Sciences, Nanjing Normal University, Nanjing, China; Department of Life Science, National Taiwan Normal University, Taipei, China

**Keywords:** Mitochondrial genome, Lepidoptera, Papilionoidea, *Troides aeacus*, Molecular phylogeny

## Abstract

**Background:** The Lepidoptera is one of the largest insect orders. Previous studies on the evolution of Lepidoptera did not confidently place butterflies, and many relationships among superfamilies in the megadiverse clade Ditrysia remain largely uncertain. Here, we generated a molecular dataset with 78 species of lepidopterian insects, including a new complete mitochondrial genome (mitogenome) sequences of the Golden Birdwing Butterfly, *Troides aeacus,* which was listed in appendix II of CITES.

**Methods:** Based on the concatenated nucleotide sequences of 13 protein-coding genes, we constructed phylogenetic trees with Bayesian Inference (BI) and Maximum Likelihood (ML) methods, and calculated the divergence times of Lepidoptera.

**Results:** Monophyly of the Papilionoidea including skippers (Hesperiidae) is strongly supported by a high bootstrap value. Butterflies were placed sister to the remaining obtectomeran Lepidoptera, and the latter was grouped with high bootstrap supports. Additionally, Papilionidae probably diverged from the group (Hesperiidae + (Nymphalidae + Pieridae)) at 102.65 Mya, the Early Cretaceous. *T. aeacus* and the Golden kaiserihind *Teinopalpus aureus* diverged in the Cretaceous of 85.32 Mya. The age of Papilionoidea indicates that the primary break up of Gondwana may have an effect on the current distributions of butterflies.

## INTRODUCTION

The Lepidoptera is one of the largest insect orders, with more than 160,000 described species (*Kawahara and Breinholt, 2014*), and provides important model for scientific enquiry. Although many lines of evidence provide strong support for the monophyly of Lepidoptera (*Grimaldi and Engel, 2005),* relationships among superfamilies, especially those in the lower Ditrysia, remain largely uncertain. The most disputed questions in lepidopteran phylogeny is the position and monophyly of butterflies, which remains unclear (*Regier et al., 2009; Cho et al., 2011*; *Mutanen et al., 2010; Regier et al., 2013; Kawahara and Breinholt, 2014*). Morphological treatments considered butterflies as close to Geometroidea (*Kristensen and Skalski, 1998; Minet, 1991; DeJong, 1996),* and recent studies suggested that butterflies might belong to the lower ditrysian lineages, though these results were weakly supported (*Regier et al., 2009; Cho et al., 2011; Mutanen et al., 2010; Regier et al., 2013*). *Cho et al. (2011)* has suggested that butterflies might be a paraphyletic assembly, however, relationships among butterfly families are still being debated (*Heikkila, 2012). Kawahara and Breinholt (2014)* presents a transcriptome-based tree of Lepidoptera that supports monophyly of butterflies. Lepidoptera are unfortunately characterized by a lack of fossils that can be confidently assigned to extant clades (*Grimaldi and Engel, 2005*). Recently, phylogenomics analysis of the divergence times in insects (*Misof et al., 2014),* which include several lineages of the Lepidoptera, suggested that the Lepidoptera diverged in the early Cretaceous.

Papilionidae is one of the most known families within the Lepidoptera, and its members are called swallowtail butterfly, which is the best-known family and may be the most well-known group of invertebrate animals (*Simonsen et al., 2011*). Phylogenetic relationships within the Papilionidae have recently been studied based upon morphology and molecular biology. *Vane-Wright (2003)* noted that more work has gone into trying to understand the interrelationships of the 600 or so species of Papilionidae than any other family of Lepidoptera, and yet ‘‘schemes abound. But we remain far from any consensus’’.

The genus *Troides* is one of three genera of birdwing butterflies with the natural beauty, other two genera of which are *Ornithoptera* and *Trogonoptera* (*Sadaharu et al., 1999*). It is supposed that their ancestors have divergened either, as Gondwanaland began to fragment, in the region that includes Australia and New Guinea Island, or in a part of South-East Asia (*Ridd, 1971; Talring, 1972; Sasajima et al., 1978; Audrey-Charles, 1983*). The Golden Birdwing butterfly, *Troides aeacus,* is a member of the genus *Troides* within the family Papilionidae (*Kristensen and Skalski, 1999). T. aeacus* is a beautiful and large butterfly, distributed locally in northern India, Nepal, Burma, China, Thailand, Laos, Vietnam, Taiwan, Cambodia, peninsular Malaysia and Indonesia (*Chou, 1998; Kunte, 2000; Wu, 2002*). This species is the northernmost birdwing butterfly as it stretches its range well into China. It is divided into five subspecies distributed throughout tropical southeastern Asia (Wu *et al*., 2010). The subspecies *T. aeacus aeacus* Felder distributes in southern China, most parts of the Indo-chinese Peninsula and northeastern India; *T. aeacus formosanus* (Rothschild) is endemic to Taiwan; *T. aeacus szechwanus* Okano & Okano occurs in the western China; the little known *T. aeacus insularis* Ney is restricted to Sumatra; and *T. aeacus malaiianus* Furhsorfer is confined to the Malayan Peninsula (Wu *et al*., 2010). Li et al. (2010) summarized the biology and habitat of *T. aeacus* with GLM, and found that its abundance will increase with both the number of adult nectar plants and larval host plants, while it will decrease with dense forest canopy.

Therefore, they developed first recommendations to conserve this species (*Li et al., 2010*). As all *Troides* species are included in appendix II of the Convention on International Trade in Endangered Species of Wild Life and Flora (CITES) (*Collins and Morris, 1985; CITES, 2008),* understanding of the structure and function of the genome of *T. aeacus* has considerable implications on researches of conservation genetics of these rare butterflies. However, despite the significance of *Troides* as a conservation species, only limited mitochondrial gene sequences are available. For instance, *Yuan (2003)* have done the molecular taxonomy on the genus *Troides* from China based on the ND5 and *CoxII* sequences; *Tsao and Yeh (2008)* presented partial *Cox1* sequence of *T. aeacus kaguya.* Nevertheless, the phylogenetic position and divergence times of *T. aeacus* are still unclear.

In the present study, we sequenced the complete mitogenome of *T. aeacus.* The aims of the present study were to develop molecular markers for phylogenetics, identification and species delimitation within the genus *Troides*, to estimate divergence time of this species, and to examine the evolution in Lepidotera. Phylogenetic analyses among insects were done, and the utility of mitogenome for lepidopteran phylogeny construction was briefly discussed. The sequences given in this study not only may provide information useful for phylogenetics of Lepidoptera and other insects, but also develop genetic markers for species identification in the *Troides* species complex. The complete mitogenome sequence was compared with mitogenomes of the other insect orders, and features distinctive from other insects were reported herein. In addition, we used our mitogenome data to estimate divergence times of the major lineages within the Lepidoptera with a Bayesian relaxed clock method (*Drummond et al., 2006*).

## Materials & Methods

### Biological materials

A *T. Aeacus* adult was collected from Xishuangbana, Yunnan Province in China. After examination of the external morphology for the identification of *T. aeacus*, the genomic DNA was extracted from the legs of an adult specimen by using proteinase K/SDS method. The DNA samples were stored at -20°C and used as a template for subsequence PCR reactions.

### Laboratory procedures

Some partial sequences of *ND2-COI, COI, COII, ND5, Cytb* and *12S* were amplified and sequenced at first using primers (Supplementary Table S1). These fragments of *T. aeacus* mitogenome were amplified by using Takara rTaq^TM^ (Takara Bio, Otsu, Shiga, Japan). Based on these sequences information, some new pairs of primers (*COI-COII-F & COI-COII-R, COII-ND5-F & COII-ND5-R, ND5-Cytb-F & ND5-Cytb-R, Cytb-12S-F & Cytb-12S-R)* were designed which of them each segment overlapped the adjacent sequence by 80-150 bp to amplify the larger mtDNA fragments of *T. aeacus* using Takara LA Taq^TM^ (Takara Bio, Otsu, Shiga, Japan) by long PCR, and the products were tested by electrophoresis on an agar gel. Actually another primer pair (*12S-ND2-F & 12S-ND2-R)* was designed aimed to amplify the A+T-rich region and the fragment was ligated into pMD19-T vector (TaKaRa Bio, Otsu, Shiga, Japan). All of them were stained with ethidium bromide, and photographed under ultraviolet light.

All fragments were sequenced in both directions, and large PCR products were amplified by primer walking strategy. Sequences were checked and assembled by using DNASTAR Lasergene software (DNASTAR, Inc., Madison, WA, USA), BioEdit 7.0.9 (*Hall, 1999)* and Chromas v2.22 (http://www.technelysium.com.au/chromas.html). Then we checked manually to obtain the complete mitogenome sequence of *T. aeacus.*

### Data analyses

Protein-coding genes were identified by using SEQUIN (Ver 5.35) (http://www.ncbi.nlm.nih.gov/projects/Sequin/download/seq_win_download.html) and compared with the corresponding known complete mtDNA sequences of Lepidoptera and other related insects. 22 *tRNA* genes were identified using software tRNAscan-SE 1.21 (*Lowe and Chan, 2016)* and their cloverleaf secondary structure and anticodon sequences were identified using software DNASIS ver. 3.6 (Hitachi Software Engineering Co., Ltd).

### Phylogenetic analyses

A total of 78 examined species representing 68 genera of 18 families of 9 superfamilies were chosen for the phylogenetic analyses. The GenBank accession numbers used in this study are listed in Supplementary Table S2. *Paracladura trichoptera, Jellisonia amadoi, Haematobia irritans irritans* and *Bittacus pilicornis* were chosen as outgroups (Supplementary Table S3). DNA alignment was inferred from the nucleic acid alignment of each of the 13 protein-coding genes by using MEGA 7.0 (*Kumar et al., 2016*). MrModeltest 2.3 (*Nylander, 2004)* was used to select the model for the Bayesian Inference (BI) and Maximum Likehood (ML) analyses, respectively. We created thirteen datasets to respectively calculate the best model for each protein-coding gene. According to the Akaike information criterion, the GTR + G model was selected as the most model appropriate for ATP8. Other datasets were used GTR+I+G model. The BI analysis was performed using MrBayes vers. 3.1.2 (*Ronquist and Huelsenbeck, 2003*) under both of the models. The analysis was run twice simultaneously for 10,000,000 generations with every 1000 trees sampled. We discarded the first 1000 000 generations (1000 samples) as burn-in (based on visual inspection of the convergence and stability of the log likelihood values of the two independent runs). The Maximum likelihood (ML) analysis was performed using the program RAxML ver. 8.0 (*Stamatakis, 2014*) with the same model. A bootstrap analysis was performed with 1000 replicates. Resulting tree files were inspected in FigTree v1.4.2 (http:// tree.bio.ed.ac.uk /software/figtree/).

### Divergence time estimation

Analyses were based on sequences of 13 protein coding genes from 78 taxa sampled, include four outgroups. The program BEAST v 2.1.3 (*Bouckaert et al., 2014*) was used to estimate times of divergence, with calibrations using five nodes (Supplementary Table S4). We used the result from Wahlberg et al. as secondary calibration point to calibrate the age of the first split in Nymphalidae at 90 Mya (*Wahlberg et al., 2009*) and Papilionoidea at 104 Mya (*Wahlberg et al., 2013*). According to the result of *Zakharov et al. (2004),* the estimated age of Papilionidae and the subfamily Papilioninae was approximately 26±7 and 16.8±2.7 MY, respectively. The age of *Papilio machaon* was calibrated using 0.56±0.77 MY. Finally, we used the result from *Misof (2014)* to calibrate the root of Lepidoptera.

In our study, the Bayesian relaxed clock analyses were carried out with the program BEAST v2.2.3 (*Bouckaert et al., 2014*). The XML file for the beast analysis was created using BEAUti (in the BEAST package) with the following non-default settings and priors: the site model was set to the GTR+Γ distribution with default parameters, the clock model was set to a relaxed clock with uncorrelated rates, and the tree model was set to a Yule process of speciation. The Markov chain Monte Carlo (MCMC) analyses were run for 100 million generations, sampling every 2000 generations and the first 25% discarded as burn-in. We used Tracer v1.5 to assess whether the likelihood traces of the four runs had converged to a stable equilibrium and that ESS values were above 200 for all parameters.

## Results

### *Troides aeacus* mitogenome

The organization of the *T. aeacus* mitogenome is shown in *Fig. 1.* The complete mitogenome of *T. aeacus* is a circular molecule 15,341 bp in size and has been deposited in GenBank (Accession number: EU625344). Each of the 37 genes found in a typical metazoan mitogenome was present: 13 protein coding genes (*ATP6, ATP8, COI-III, ND1-6, ND4L and Cytb),* 2 ribosomal *RNAs (12S rRNA* and *16S* rRNA), 22 transfer *RNAs,* and a putative control region (A+T rich region) (*Supplementary Table S5*).

**Figure 1.**
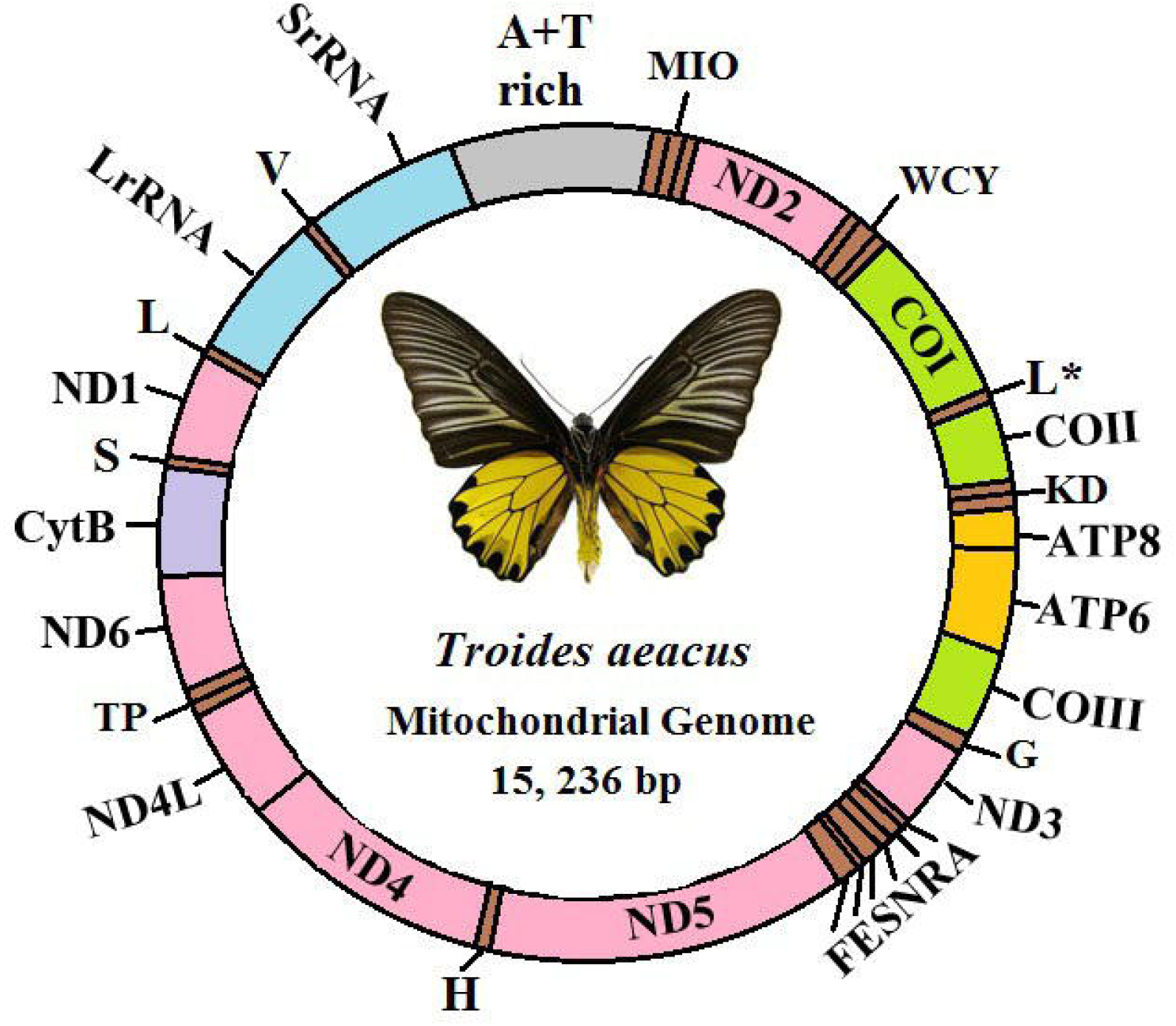
Circular map of the T *aeacus* mitogenome. The abbreviations for the genes are as follows: *COI-III* refer to the cytochrome oxidase subunits, *CytB* refers to cytochrome *b,* and *ND1–6* refer to *NADH* dehydrogenase components. tRNAs are denoted as one-letter symbol according to the IUPAC-IUB single letter amino acid codes.

We calculated all the base compositions of the mtDNA using DNASTAR (DNASTAR Inc., USA). The average C+G content of the *Troides* mitogenome is 19.7%, and the average A+T content 80.3%. This corresponds well to the AT bias generally observed in insect mitogenome, which ranges from 69.5% to 84.9% (*Crozier and Crozier, 1993; Dotson and Beard, 2001*). The coding of the mitogenome is very compact and some of the genes overlap each other in the mitogenome. The *T. aeacus* mitogenome contains 22 *tRNA* genes, which are interspersed in the genome and range in size from 63 to 134 bp in length. They were identified in the same relative genomic positions as observed for the *Coreana raphaelis* mitogenome (*Kim et al., 2006*). All of them showing typical cloverleaf secondary structures and their anticodons are similar to those found in other metazoan animals (*Supplementary Fig. S1*). Overlapping was observed for *tRNA^Trp^/tRNA^Cys^,* as reported by *Lessinger (2000),* producing separate transcripts from their opposite directions.

In insects, common Met start codons (ATA or ATG) could be assigned to nine of the protein coding genes. In the *T. aeacus* mitogenome all protein-coding genes start with an ATG codon except for the *ATP8, ND3, ND5, ND4, ND4L* and *ND6.* The *ATP8, COII, ND6, ND1* and *Cytb* genes are terminated with TAA. Among these 13 protein-coding genes, the *COI, COII, ATP6, ND5, ND4L* and *ND4* genes are ones that adjoin at their 3’-end region to another gene. The *COI, COII, ND5* and *ND4* all end with *T-tRNA, ATP6* ends with *T-COIII,* and *ND4L* stops with *TA-ND4.* The *COI* gene must use a nonconventional start site because no regular start codon is available after the last stop codon upstream to the open reading frame (ORF) of *COI.* There is 4-bp putative initiation codon, TTAG, before the CGA codon as suggested in the genus *Bombyx* (*Yukuhiro et al., 2002),* and the *T. aeacus COI* gene also starts with TTAG and ends with T-tRNA. Similar results were obtained in other insects like *Adoxophyes honmai* (*Lee et al., 2006),* and the use of nonstandard translational start sites for *COI* has also been proposed for many other insect species (*Lessinger et al., 2000; Yukuhiro et al., 2002*). More studies for mRNA transcripts would be needed to clarify prediction on the position of *COI* initiation (*Lee et al., 2006*). The *COI* gene has been described for other insect species like *A. honmai* where it has been proposed that the termination codon can be generated by polyadenylation (*Lee et al., 2006*).

The non-coding regions include the control region and a few spacers. There are 25 non-coding regions found in the *T. aeacus* mitogenome. There is a *tRNA^Glu^* gene within the ŕRNA^Ser^gene, which is very different from *Coreana raphaelis* (*Kim et al., 2006*). The *tRNA^Glu^* and *tRNA^Ser^* are out of the picture in the *C. raphaelis* mitogenome. The *T. aeacus* mitogenome possesses a total of 111 bp intergenic spacer sequences and these are spread over 12 *protein-coding* regions. The longest one is an 37 bp long spacer sequence located between the *tRNA^Leu^* and *16S rRNA* gene, containing a high A+T nucleotide (89.1%).

The A+T-rich region is higher in variation and there are also some repeats in this region (*Moritz, 1987*). It contains several short repeating sequences (6–11 bp) with varying copy number scattered through the whole A+T-rich region of *T. aeacus.* This repeat unit occurs seven times and is 92 bp in total length (*SupplementaryFig. S2*).

### Phylogenetic analyses

Different optimality criteria and dataset compilation techniques have been applied to find the best method of analyzing complex mitogenomic data (*Castro and Dowton, 2005; Kim et al., 2006; Stewart and Beckenbach 2005*). We combined data from the other insect mitogenomes with the *T. aeacus* mitogenome sequences to gain some insights on the phylogeny of Lepidoptera using the combined mitochondrial protein alignments (*Foster and Hickey, 1999*). To further elucidate the phylogenetic relationships among 78 species of insects, we constructed phylogenetic trees based on the concatenated sequences of 13 protein coding genes (*Figs. 2, 3*). For rooting we used the corresponding sequences of *Paracladura trichoptera, Jellisonia amadoi, Haematobia irritans irritans* and *Bittacus pilicornis* as outgroups.

**Figure 2.**
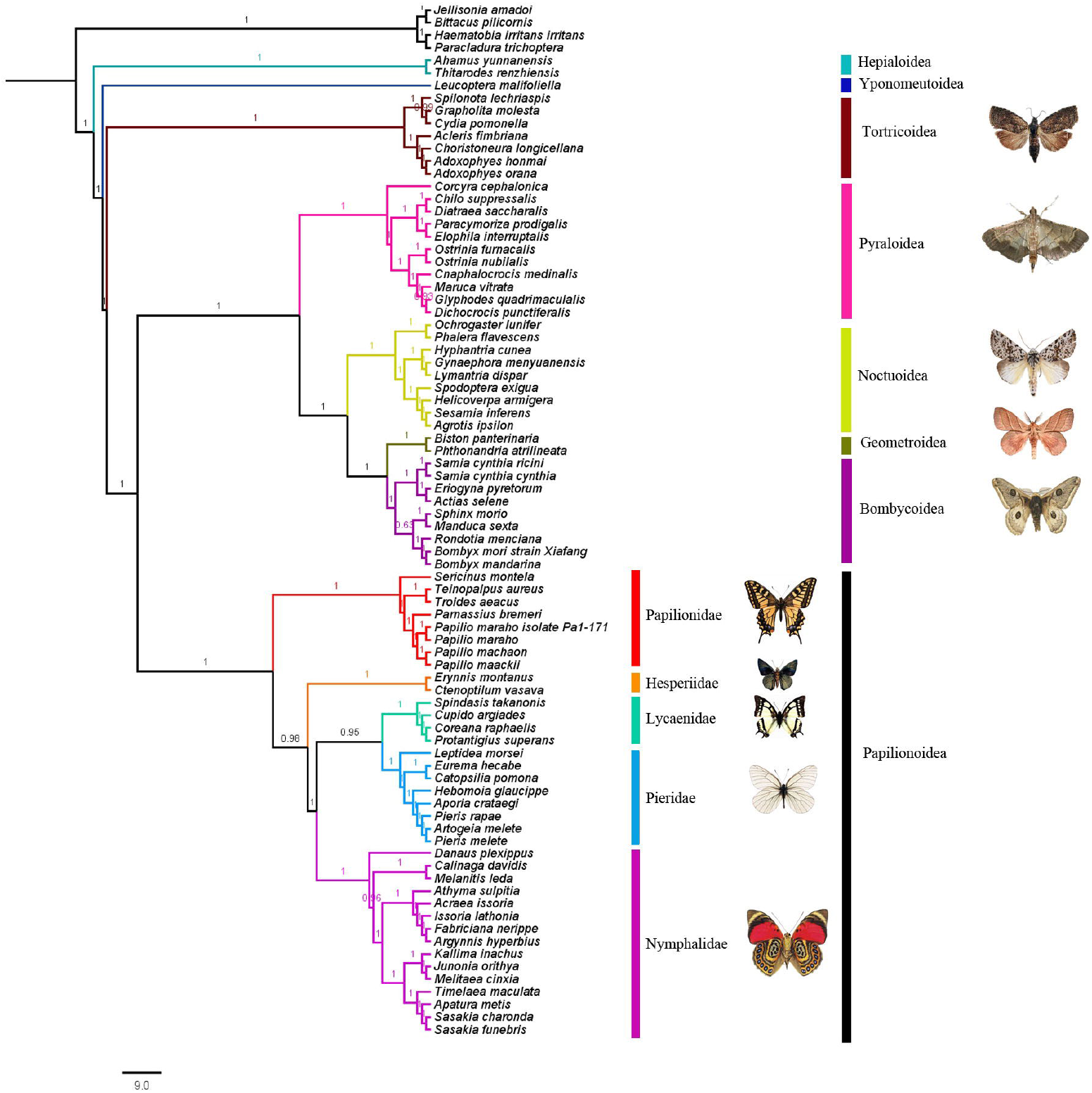
Phylogenetic tree estimation from the mitogenome sequences of selected insects using Bayesian Inference (BI) method. Numbers at each branch are bootstrap values.

**Figure 3.**
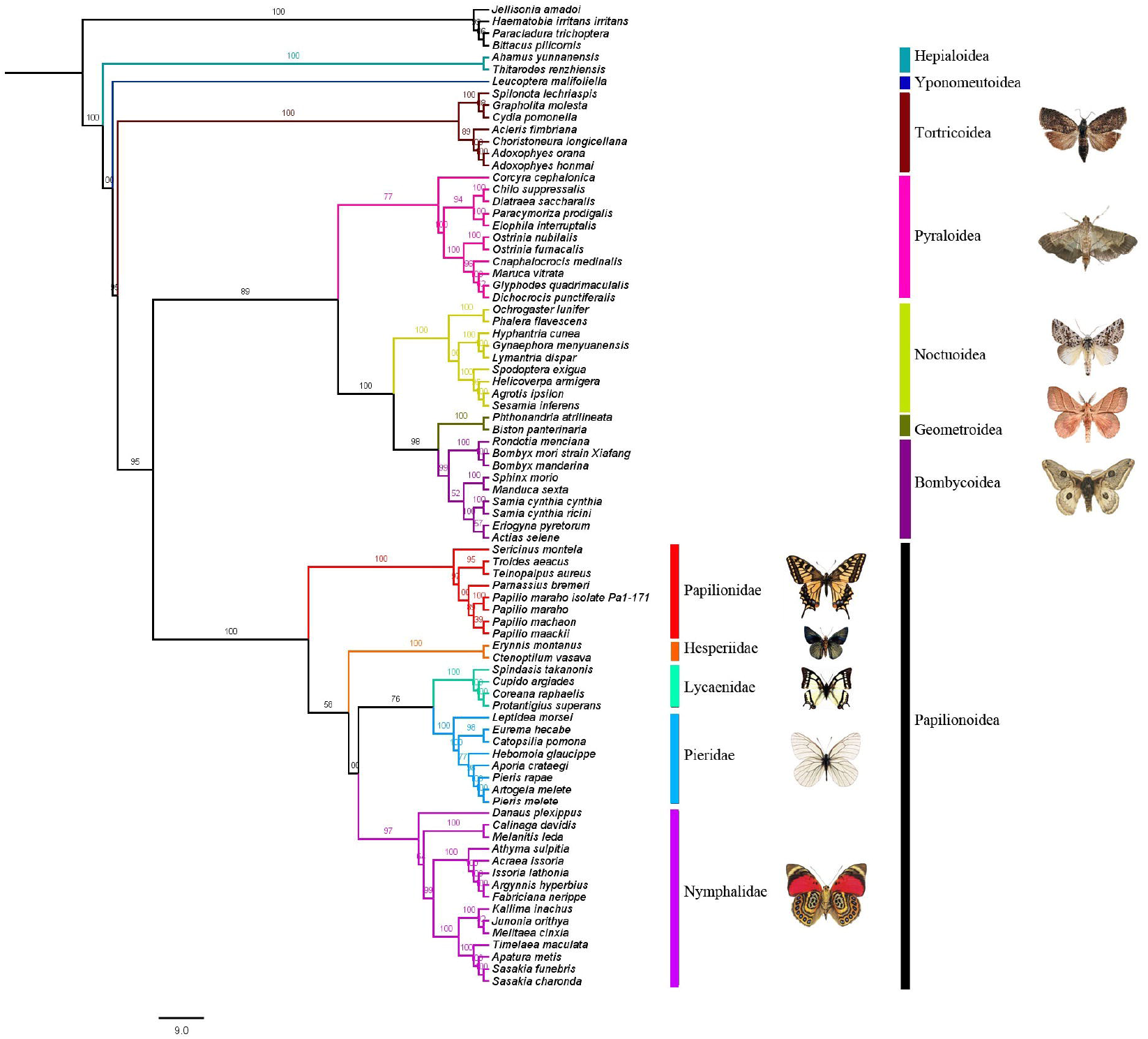
Phylogenetic tree estimation from the mitogenome sequences of selected insects using Maximum Likehood (ML) method. Numbers at each branch are bootstrap values.

ML and BI analyses recovered similar topologies for all individual-marker analyses. The two trees differed in the resolution of some nodes, but the two methods never strongly supported conflicting topologies. All groupings were highly supported, and the results were consistent with previous reports (*Kawahara and Breinholt, 2014; Wahlberg et al., 2013*). All analyses recovered Tortricoidea, Hepialoidea, Yponomeutoidea, Bombycoidea, Geometroidea, Noctuoidea, Pyraloidea and Papilionoidea as monophyletic groups. The Pyraloidea are confidently placed as the sister group to Noctuoidea + Geometroidea + Bombycoidea (100% BP), which were placed at the sister group to butterflies. Within the Papilionoidea, all nodes are supported by more than 80% bootstrap supports. The Papilionidae is placed as sister to (Hesperiidae + (Nymphalidae + (Lycaenidae + Pieridae))). Our BI and ML analysis results support the idea that Noctuoidea is sister to (Geometroidea + Bombycoidea).

With regard to relationships among the families within the Papilionoidea, the (Pieridae + Lycaenidae) clade is sister to the Nymphalidae. This result is different with a recent analysis of Lepidoptera which used more extensive molecular data (*Regier et al., 2013*). They placed Pieridae as the sister to (Lycaenidae + Nymphalidae). Within the Papilionidae, Leptocircini and Troidini were sister groups, with Papilionini being sister, this result is different from the results of *Miller (1987)* and *Tyler et al. (1994).* The placement of *T. aeacus* is placed as sister to *Teinopalpus aureus* in the Papilionidae clade.

### Divergence time estimation

The estimated divergence times among the Lepidoptera are shown in *Fig. 4*. The first divergence in Lepidoptera happened in the late Jurassic (about 140 Mya), and most clades have been diverged throughout the Cretaceous (about 120 Mya). As showed in previous molecular studies (*Wahlberg et al., 2013; Heikkilä et al., 2012),* Papilionidae was diverged from the group (Hesperiidae + (Nymphalidae+ Pieridae)) in the Cretaceous, between 110 and 90 Mya. Interestingly, in this study, the clade Pyraloidea and the huge clade of moths including Noctuoidea, Geometroidea and Bombycoidea, be nearly the same age as the butterflies, about 100 million years. This result is consistent with a recent study (*Wahlberg et al., 2013*). Lineages leading to extant families all diverged from 102.65 Mya, with Papilionidae diverging from the common ancestor to the other butterflies at about 91 Mya, Hesperiidae about 84.5 Mya, Nymphalidae about 92 Mya, Lycaenidae about 74 Mya, and Pieridae about 83 Mya.

**Figure 4.**
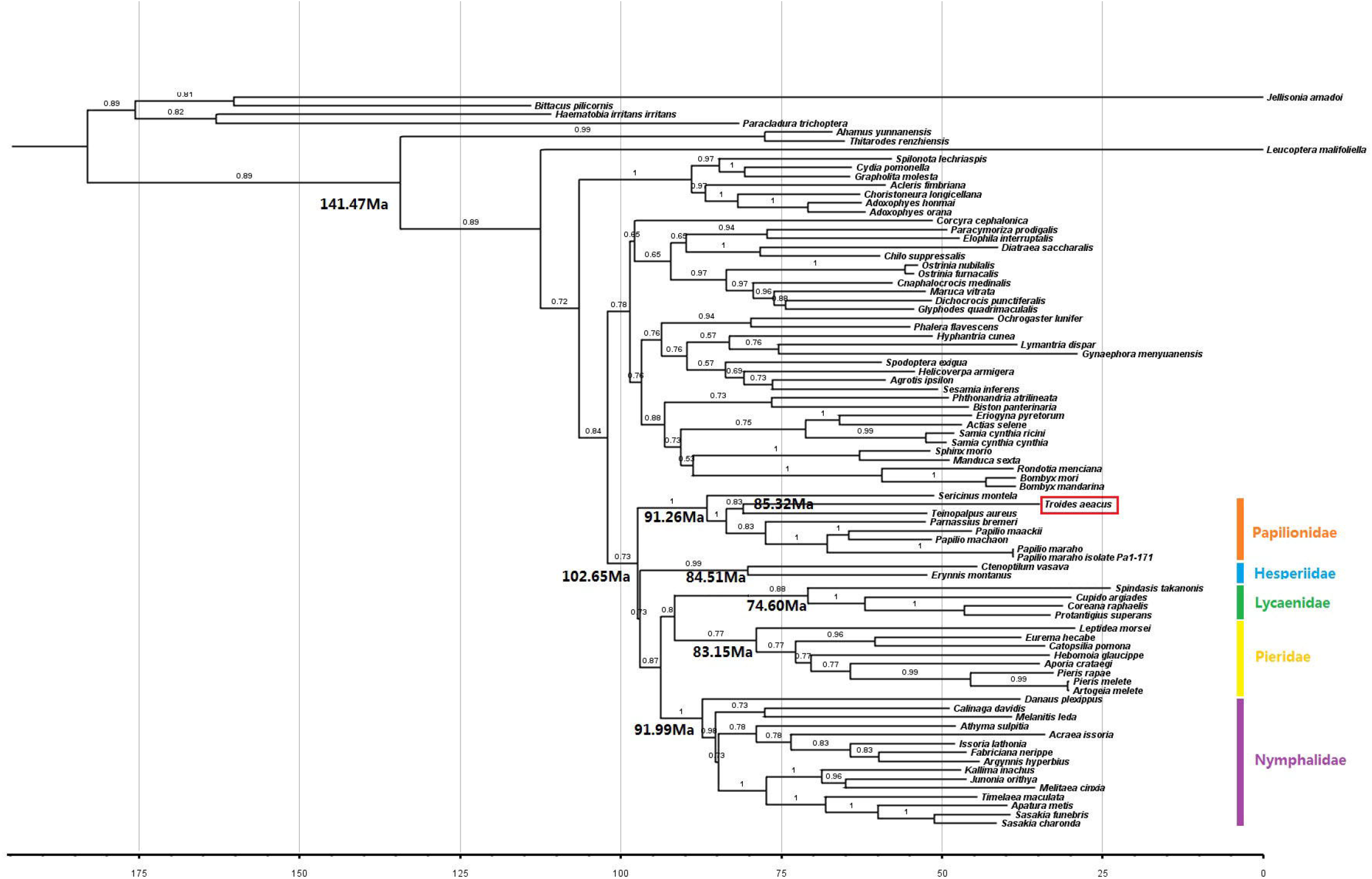
Estimated times of divergence for lineages leading to the families of Lepidoptera.

## Discussion

### *Troides aeacus* mitogenome

The size of *T. aeacus* mitogenome is longer than the mitogenome sizes reported for the other butterflies which range from 15,140 bp in *Artogeia melete* (Pieridae) (*Kim et al., 2006*) to 15,314 bp in *Coreana raphaelis* (Lycaenidae)(Hong *et al*., 2009). The gene order and orientation of the mitogenome are identical to that of *Drosophila yakuba* (*Clary and Wolstenholme, 1987),* the structure of which is conserved as in the divergent insect orders and even some crustaceans (*Crease, 1999; Nardi et al., 2003*). Such an extensive conservation in orientation and gene order in divergent insects has been inferred to be ancestral for insects (*Boore et al., 1998*). The arrangement of mitochondrial coding genes in *T. aeacus* matches those of many distantly related insects, showing that *T. aeacus* shares the mitogenome organization considered ancestral to insects and crustaceans (*Hwang et al., 2001*).

### Phylogenetic relationships

In conclusion, our phylogenetic analyses suggest the moth superfamilies Tortricoidea, Hepialoidea, Yponomeutoidea, Bombycoidea, Geometroidea, Noctuoidea and Pyraloidea originally defined are monophyletic. It is important that our result conforms to internal relationships among taxa within the Insecta in the traditional classification system. Besides, protein-coding genes of mitogenome are valuable to the studies of speciation and evolution of species in the Cambrian (*Curole and Kocher, 1999; Hwang et al., 2001*).

For the superfamilies Hepialoidea, Yponomeutoidea and Tortricoidea, our results are similar to the study in *Mutanen et al. (2010),* which based on combined analysis from the mitochondrial and nuclear gene data. Interestingly, in previous studies (*Kristensen, 2003; Cho et al., 2011),* there were no consistent evolutionary relationships among Noctuoidea, Bombycoidea and Geometroidea. *Kristensen (2003)* suggested that the group (Geometroidea and Hesperioidea) is sister to the group (Noctuoidea and Bombycoidea). However, *Cho et al. (2011)* proposed a sister-group relationship for the Geometroidea and (Noctuoidea + Bombycoidea).

Prior molecular phylogenetic work suggested that butterflies were more closely related to ‘microlepidoptera’ than to the large moths, but none of these recent studies could confidently confirm this conclusion (*Regier et al., 2009; Cho et al., 2011; Mutanen et al., 2010; Regier et al., 2013*).

According to *Minet (1991),* Papilionoidea are closely related to Geometroidea, whereas according to *Regier et al. (2009),* they recovered Bombycoidea as the sister taxon of Noctuoidea. Our results consistent with the relationships revealed by them. *Kawahara and Breinholt (2014)* placed Bombycoidea as the sister of (Geometroidea + Noctuoidea). However, our results show that the butterfly clade is well supported and is placed sister to the clade (((Bombycoidea + Geometroidea) + Noctuoidea) + Pyraloidea).

Papilionoidea are well supported as monophyletic. Papilionidae, Lycaenidae, Hesperiidae and Pieridae are strongly supported as monophyletic group. The relationships within Nymphalidae recovered are consistent with the recent molecular analyses (*Nélida et al., 2009),* which groups Satyrinae as sister to the clade (Nymphalinae + (Limenitidinae + Heliconiinae)) and, the Danainae as the basal subfamily, sister to the clade (Satyrinae + (Nymphalinae + (Limenitidinae + Heliconiinae))), and this result is distinct from some previous studies (*Regier et al., 2013; Mutanen et al., 2010; Cho et al., 2011*).

Within the Papilionoidea, we recovered all families to the traditional classification. The placement of Hesperiidae is consistent with previous studies (*Kawahara and Breinholt, 2014; Wahlberg et al., 2013),* which are placed on the clade (Papilionidae + (Hesperiidae + (Lycaenidae + Nymphalidae))), whereas it is contrary to *Heikkilä’s (2012)* hypothesis about the position of Papilionidae as sister-group to Hesperiidae.

### Divergence time estimation

*Vane-Wright (2004)* highlighted that our knowledge of the age of butterflies is going through a period of “adolescence”, and that future studies would help us get more insight into how long butterflies have existed. According to the *Misof et al. (2014),* they dated the Lepidoptera to the Early Cretaceous, contemporary with the radiation of flowering plants. The estimate of 91.26 Mya for the origin of the family Papilionidae is much older than the minimum age of 48 Mya indicated by fossils (*Zakharov et al., 2004*). However, it is substantially early than the result of *Condamine (2013),* in which swallowtails was recovered in the Eocene c. 52 Mya (CI 46–62.5 Mya) around the early Eocene Climatic Optimum. It may be because the fossil calibrations used represented minimum age constraints. In addition, they used a maximum age of 183 Mya to set the root, based on the assumption that all but the earliest clades of the order Lepidoptera diversified in association with angiosperm plants (*Condamine et al., 2013*).

The divergence times for the major clades within Papilionidae suggested in this study are corresponding roughly to the divergence times for the major clades in the Nymphalidae, and are considerably older than the Tertiary suggested by *Simonsen et al. (2011)* and *Wahlberg (2006). Wahlberg et al. (2013)* suggested that Lepidoptera underwent a radiation in the Late Cretaceous at ca. 90 Mya. But our result is much older, up to 102.65 Mya. Thus, the age of Papilionoidea indicates that the primary break up of Gondwana (i.e. the splitting of Africa from Gondwana about 100 Mya) may have an affect on the current distributions of butterflies.

## Conclusions

The organization of the *T. aeacus* mitogenome is identical with that of the general invertebrates. It is rich in A+T throughout the whole mitogenome, and should be used for developing mitogenome genetic markers for species identification of other *Troides* species complex, which is listed in the appendix II of CITES species. Thus, our results should have important implications on the research of conservation genetics for these precious and endangered birdwing butterflies. Sequences from mtDNA and other genetic markers have great potential for clarifying the genetic relationships among Insecta, and mitogenome data presented here may promote efforts to resolve taxonomic problems in Insecta (*Stewart and Beckenbach, 2005*).

Our result is consistent with previous studies that strongly contradict historical placement of butterflies. The placement of Hesperioidea in this study is contrary to Heikkilä’s hypothesis of the position of Papilionidae as sister-group to Hesperiidae. In addition, as in previous molecular studies, Papilionoidea probably diverged in 102.65 Mya, the Early Cretaceous.

According to our results, the extant diversity of Lepidoptera has evolved since the Lower Cretaceous, an important period for the diversification.

## Acknowledgements

We thank Dr. S. X. Xu, Dr. Y. Xiong, N. X. Wang and Y. P. Yao (College of Life Sciences, Nanjing Normal University) and Profs. Jia-Sheng Hao, Hong-Chun Pang,,Dr. L. Wei, S. Y. Cao, X. B. Zuo (College of Life Sciences, Anhui Normal University) for valuable helps with experimental analyses.

## Additional information and declarations

### Author Contributions

- Si-Yu Dong performed the experiments, analyzed the data, wrote the paper, prepared figures and/or tables.
- Guo-Fang Jiang conceived and designed the experiments, analyzed the data, wrote the paper, prepared figures and/or tables, reviewed drafts of the paper, expertise on study object, field work.
- Gang Liu performed the experiments, field work, and wrote the paper.
- Fang Hong prepared figures and/or tables, performed most of the manuscript revision.
- Yu-Feng Hsu reviewed drafts of the paper.

## Supplemental Information

Supplementary data [Fig.S1-S2;Tables S1-5] associated with this article can be found online at

