## Supplemental Table S1 for "The complete mitochondrial genome of the Golden Birdwing butterfly *Troides aeacus*: insights into phylogeny and divergence times of the superfamily Papilionoidea"

**Table S1. List of PCR primers used in this study**

| Name Direction | Sequence |
| --- | --- |
| ND2-COI F  ND2-COI R  COI F  COI R  COI-COII F  COI-COII R  COII F  COII R  COII-ND5 F  COII-ND5 R  ND5 F  ND5 R  ND5-Cytb F  ND5-Cytb R  Cytb F  Cytb R  Cytb-12S F  Cytb-12S R  12S F  12S R  12S-ND2 F  12S-ND2 R | TGGAGTTTTAGGAGGTATT  CTAGAAATGGAGGAAGTCCTC  TTTCTACAAATCATAAAGATATTGG  TAAACTTCAGGGTGACCAAAAAATCA  CCCTATTTGTCTGAGCTGTAGGAA  AATATAACCTACTAAAATAGTAA  GAGACCATTACTTGCTTTCAGTCACT  CTAATATGGCAGATTATATGTATTGGA  GATGTTGATAATCGTATTGTT  CCTCTTACTACTTTATGTT  GCTAATTATGAATTTGATT  GATACTCTTCATCATATA  CCTCCTATATAACGAATATC  AGTAGCATTATCAATAGCA  TACGTTTTACCATGAGGTCAAATATC  ACTTCTTTTCTTATGTTTTCAAAAC  CCGACCTGTTGAAGATCCTTAT  TCAGATCAAGATGCCGATT  AAGAGCGACCGGCGATGTGT  AAACTAGGATTAGATACCCTATTAT  TCTAGGAACACTTTCCAGT  CTAAGAAAGGGGGTAAACCTC |

F indicates Forward and R refers Reverse primers, respectively.
