## Supplemental Table S2 for "The complete mitochondrial genome of the Golden Birdwing butterfly *Troides aeacus*: insights into phylogeny and divergence times of the superfamily Papilionoidea"

**Table S2. Accession numbers of complete mitogenome for Lepidoptera retrieved from Genbank.**

| **Superfamily / Family** | | **Species** | **Acc. number:** |
| --- | --- | --- | --- |
| Tortricoidea | Tortricidae | *Cydia pomonella* | NC_020003.2 |
|  |  | *Grapholita molesta* | NC_014806.1 |
|  |  | *Spilonota lechriaspis* | NC_014294.1 |
|  |  | *Adoxophyes orana* | NC_021396.1 |
|  |  | *Adoxophyes honmai* | NC_008141 |
|  |  | *Choristoneura longicellana* | NC_019996.1 |
|  |  | *Acleris fimbriana* | NC_018754.1 |
| Hepialoidea | Hepialidae | *Ahamus yunnanensis* | NC_018095.1 |
|  |  | *Thitarodes renzhiensis* | NC_018094.1 |
| Yponomeutoidea | Lyonetiidae | *Leucoptera malifoliella* | NC_018547.1 |
| Bombycoidea | Bombycidae | *Rondotia menciana* | NC_021962.1 |
|  |  | *Bombyx mori strain Xiafang* | AY048187.1 |
|  |  | *Bombyx mandarina* | AB070263.1 |
|  | Sphingidae | *Sphinx morio* | NC_020780.1 |
|  |  | *Manduca sexta* | NC_010266.1 |
|  | Saturniidae | *Samia cynthia ricini* | NC_017869.1 |
|  |  | *Samia cynthia cynthia* | KC812618.1 |
|  |  | *Eriogyna pyretorum* | NC_012727.1 |
|  |  | *Actias selene* | NC_018133.1 |
| Geometroidea | Geometridae | *Biston panterinaria* | NC_020004.1 |
|  |  | *Phthonandria atrilineata* | NC_010522.1 |
| Noctuoidea | Notodontidae | *Ochrogaster lunifer* | NC_011128.1 |
|  |  | *Phalera flavescens* | NC_016067.1 |
|  | Noctuidae | *Spodoptera exigua* | NC_019622.1 |
|  |  | *Helicoverpa armigera* | NC_014668.1 |
|  |  | *Sesamia inferens* | NC_015835.1 |
|  |  | *Agrotis ipsilon* | NC_022185.1 |
|  | Arctiidae | *Hyphantria cunea* | NC_014058.1 |
|  | Lymantriidae | *Gynaephora menyuanensis* | NC_020342.1 |
|  |  | *Lymantria dispar* | NC_012893.1 |
| Pyraloidea | Pyralidae | *Corcyra cephalonica* | NC_016866.1 |
|  | Crambidae | *Cnaphalocrocis medinalis* | NC_015985.1 |
|  |  | *Maruca vitrata* | HM751150.1 |
|  |  | *Glyphodes quadrimaculalis* | KF234079.1 |
|  |  | *Dichocrocis punctiferalis* | NC_021389.1 |
|  |  | *Ostrinia furnacalis* | NC_003368.1 |
|  |  | *Ostrinia nubilalis* | NC_003367.1 |
|  |  | *Chilo suppressalis* | NC_015612.1 |
|  |  | *Diatraea saccharalis* | NC_013274.1 |
|  |  | *Paracymoriza prodigalis* | NC_020094.1 |
|  |  | *Elophila interruptalis* | NC_021756.1 |
| Papilionoidea | Papilionidae | *Sericinus montela* | HQ259122.1 |
|  |  | *Parnassius bremeri* | HM243588.1 |
|  |  | *Papilio maraho isolate Pa1-171* | FJ810212.1 |
|  |  | *Papilio maraho* | NC_014055.1 |
|  |  | *Papilio machaon* | NC_018047.1 |
|  |  | *Papilio maackii* | NC_021411.1 |
|  |  | *Teinopalpus aureus* | NC_014398.1 |
|  | Pieridae | *Leptidea morsei* | JX274648.1 |
|  |  | *Hebomoia glaucippe* | NC_021123.1 |
|  |  | *Aporia crataegi* | NC_018346.1 |
|  |  | *Pieris rapae* | NC_015895.1 |
|  |  | *Artogeia melete* | EU597124.1 |
|  |  | *Pieris melete* | NC_010568.1 |
|  |  | *Eurema hecabe* | KC257480.1 |
|  |  | *Catopsilia pomona* | JX274649.1 |
|  | Lycaenidae | *Spindasis takanonis* | NC_016018.1 |
|  |  | *Cupido argiades* | NC_020779.1 |
|  |  | *Coreana raphaelis* | NC_007976.1 |
|  |  | *Protantigius superans* | NC_016016.1 |
|  | Nymphalidae | *Danaus plexippus* | NC_021452.1 |
|  |  | *Athyma sulpitia* | NC_017744.1 |
|  |  | *Acraea issoria* | NC_013604.1 |
|  |  | *Issoria lathonia* | NC_018030.1 |
|  |  | *Fabriciana nerippe* | NC_016419.1 |
|  |  | *Argynnis hyperbius* | NC_015988.1 |
|  |  | *Kallima inachus* | NC_016196.1 |
|  |  | *Junonia orithya* | KF199862.1 |
|  |  | *Melitaea cinxia* | NC_018029.1 |
|  |  | *Timelaea maculata* | NC_021090.1 |
|  |  | *Apatura metis* | NC_015537.1 |
|  |  | *Sasakia charonda* | NC_014224.1 |
|  |  | *Sasakia funebris* | NC_022134.1 |
|  |  | *Calinaga davidis* | NC_015480.1 |
|  |  | *Melanitis leda* | NC_021370.1 |
| Hesperioidea | Hesperiidae | *Erynnis montanus* | NC_021427.1 |
|  |  | *Ctenoptilum vasava* | NC_016704.1 |
