## Supplemental Table S3 for "The complete mitochondrial genome of the Golden Birdwing butterfly *Troides aeacus*: insights into phylogeny and divergence times of the superfamily Papilionoidea"

**Table S3. Accession numbers of complete mitogenome for outgroup retrieved from Genbank.**

| **Order** | **Superfamily / Family** | | **Species** | **Acc. number:** |
| --- | --- | --- | --- | --- |
| Diptera | Trichoceroidea | Trichoceridae | *Paracladura trichoptera* | NC_016173.1 |
| Siphonaptera | Ceratophylloidea | Ceratophyllidae | *Jellisonia amadoi* | NC_022710.1 |
| Diptera | Muscidae | Muscinae | *Haematobia irritans irritans* | NC_007102.1 |
| Mecoptera |  | Bittacidae | *Bittacus pilicornis* | NC_015118.1 |
