## Supplemental Table S4 for "The complete mitochondrial genome of the Golden Birdwing butterfly *Troides aeacus*: insights into phylogeny and divergence times of the superfamily Papilionoidea"

**Table S4. Cailbration points used to estimate times of divergence in Lepidoptera.**

| Calibration | Calibration age (mya±S.D.) | Source |
| --- | --- | --- |
| Nymphalidae | 90±5 | Wahlberg et al, 2009 |
| Papilionoidea | 104±5.4 | Wahlberg et al, 2013 |
| Papilionidae | 26±7 | Zakharov et al, 2004 |
| Papilioninae | 16.8±2.7 | Zakharov et al, 2004 |
| *Papilio machaon* | 0.56±0.77 | Zakharov et al, 2004 |
