## Supplemental Table S5 for "The complete mitochondrial genome of the Golden Birdwing butterfly *Troides aeacus*: insights into phylogeny and divergence times of the superfamily Papilionoidea"

**Table S5. Summary of mitogenome of *T. aeacus*.**

| Gene | Direction | Location | Size | IGNc | Start codon | Stop  codon |
| --- | --- | --- | --- | --- | --- | --- |
| tRNA^Met^ | F | 1-77 | 77 | 1 |  |  |
| tRNA^Ile^ | F | 79-147 | 69 | -2 |  |  |
| tRNA^Glx^ | R | 146-218 | 73 | 19 |  |  |
| ND2 | F | 238-1297 | 1060 | -2 | ATC | TAA |
| tRNA^Sec^ | F | 1296-1360 | 65 | -8 |  |  |
| tRNA^Cys^ | R | 1353-1420 | 68 | 0 |  |  |
| tRNA^Tyr^ | R | 1421-1484 | 64 | 1 |  |  |
| COI | F | 1486-3017 | 1532 | 0 | ATTAGC | T- |
| tRNA^Leu^ | F | 3018-3084 | 67 | -1 |  |  |
| COII | F | 3084-3802 | 717 | -16 | ATG | TAA |
| tRNA^Lys^ | F | 3767-3837 | 71 | -1 |  |  |
| tRNA^Asp^ | F | 3837-3903 | 67 | 0 |  |  |
| ATP8 | F | 3904-4074 | 161 | -7 | ATT | TAA |
| ATP6 | F | 4068-4744 | 677 | 3 | ATG | TAA |
| COIII | F | 4748-5538 | 791 | 46 | ATG | TAA |
| tRNA^Gly^ | F | 5585-5653 | 69 | 1 |  |  |
| ND3 | F | 5655-6012 | 358 | -2 | ATA | T- |
| tRNA^Ala^ | F | 6011-6074 | 64 | -1 |  |  |
| tRNA^Arg^ | F | 6074-6140 | 67 | 0 |  |  |
| tRNA^Asx^ | F | 6141-6206 | 66 | -2 |  |  |
| tRNA^Ser^(AGN) | F | 6205-6265 | 61 | 4 |  |  |
| tRNA^Glu^ | F | 6270-6333 | 64 | -2 |  |  |
| tRNA^Phe^ | R | 6332-6398 | 67 | 3 |  |  |
| ND5 | R | 6402-8136 | 1735 | -1 | ATT | TA- |
| tRNA^His^ | R | 8136-8200 | 65 | 0 |  |  |
| ND4 | R | 8201-9540 | 1340 | -1 | ATT | TAA |
| ND4L | R | 9540-9831 | 292 | 1 | ATT | T-ND4 |
| tRNA^Thr^ | F | 9833-9898 | 66 | 0 |  |  |
| tRNA^Pro^ | R | 9899-9962 | 64 | 2 |  |  |
| ND6 | F | 9965-10498 | 534 | 1 | ATT | TAA |
| Cytb | F | 10500-11641 | 1142 | 7 | ATA | TAA |
| tRNA^Ser^ | F | 11649-11715 | 67 | -1 |  |  |
| ND1 | R | 11732-12674 | 943 | -1 | ATG | TTA |
| tRNA^Leu^ | R | 12674-12743 | 70 | 25 |  |  |
| 16S | R | 12769-14002 | 1234 | -6 |  |  |
| tRNA^Val^ | R | 13997-14059 | 63 | 2 |  |  |
| 12S | R | 14062-14845 | 784 | -1 |  |  |
| D-loop | F | 14845-15263 | 419 |  |  |  |
