## Supplemental Fig.S1 for "The complete mitochondrial genome of the Golden Birdwing butterfly *Troides aeacus*: insights into phylogeny and divergence times of the superfamily Papilionoidea"

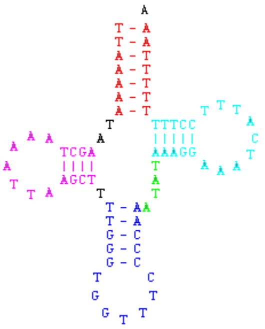
tRNA-Met

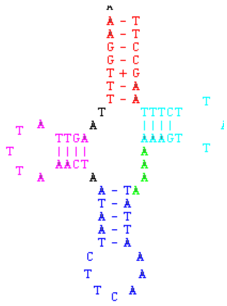
tRNA-Sec

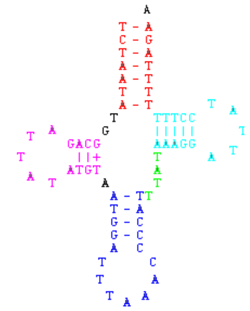
tRNA-Leu(UUN)

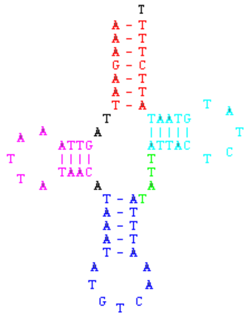
tRNA-Asp

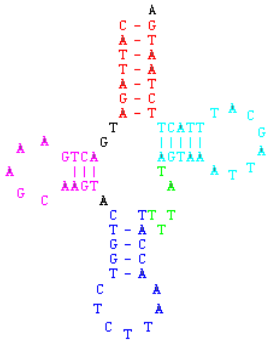
tRNA-Lys

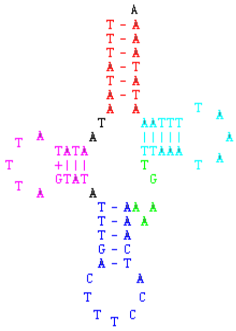
tRNA-Gly

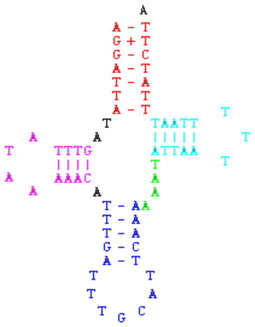
tRNA-Ala

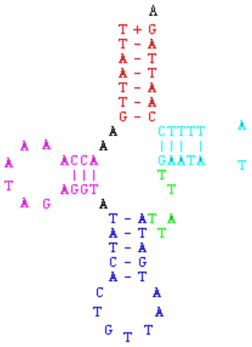
tRNA-Asn

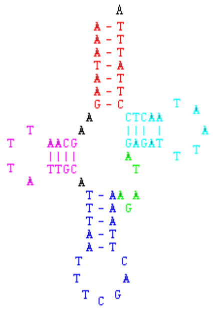
tRNA-Arg

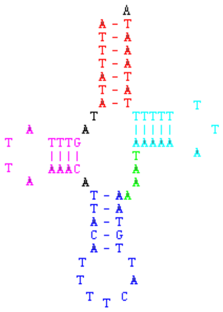
tRNA-Glu

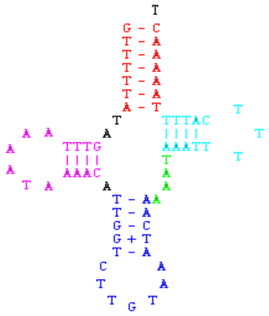
tRNA-Thr

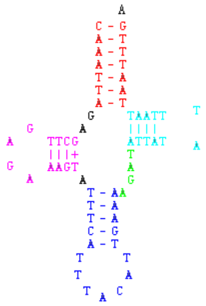
tRNA-Val

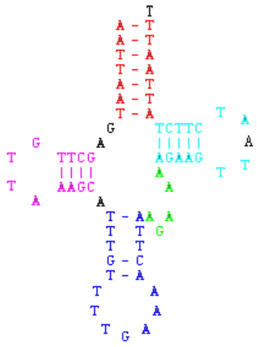
tRNA-Ser

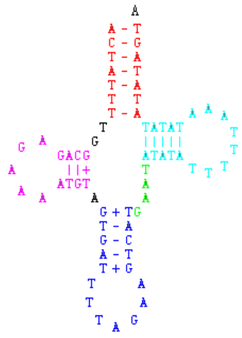
tRNA-Leu

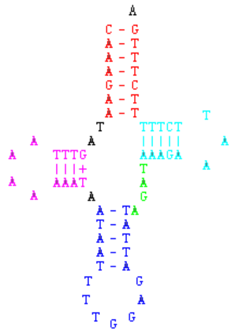
tRNA-Pro

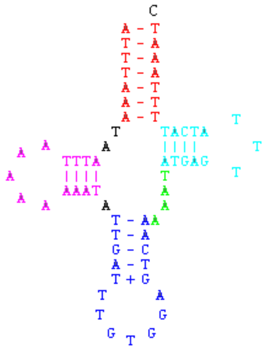
tRNA-His

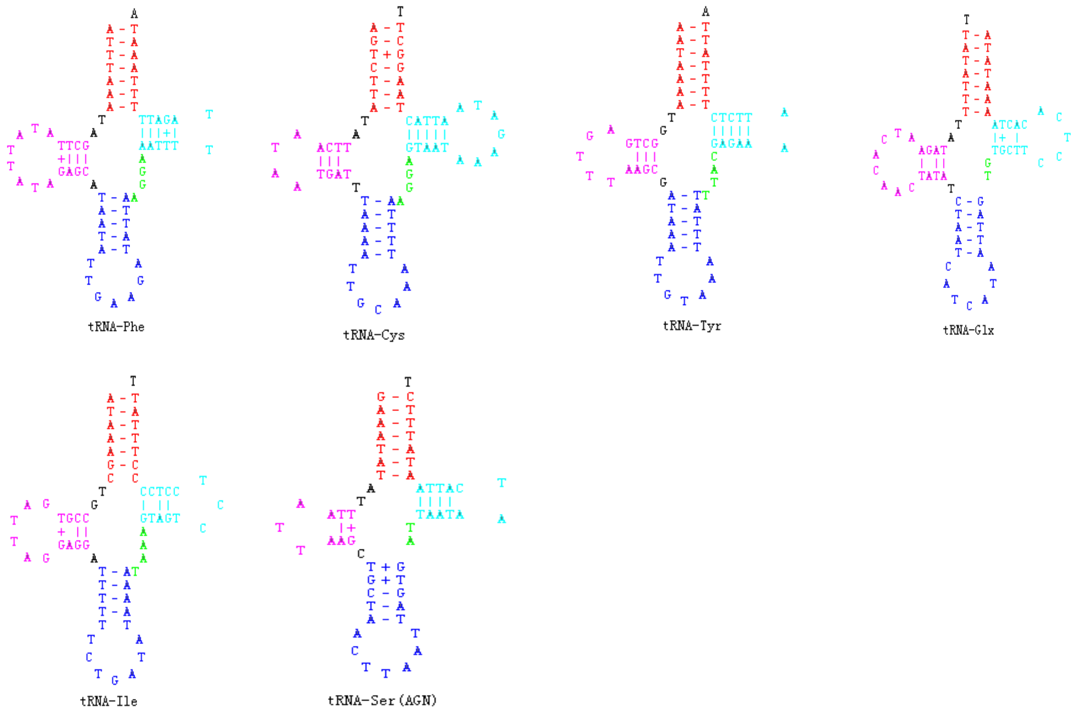


**Fig.S1. Predicated secondary clover-leaf structure for the 22 *tRNA* genes of *T. aeacus*.**
