## Supplemental Fig.S2 for "The complete mitochondrial genome of the Golden Birdwing butterfly *Troides aeacus*: insights into phylogeny and divergence times of the superfamily Papilionoidea"

ATACAAAATTTTTTCATATAGATTTTTTTTTTTTTTTTATAAATAA

AATATTTTATTATAAATTATTAAATTTTATATATTCTCTTTCTTTATC

TCTACTAATAGGGAATTAAATAATTTTTAATAAATAGAAAATTTTT

TTTATATTAAATGTTAAAAATTTAATTAATTAATATTATTATTTATTT

ATTAAATTCATTATTTAATATATTATATATATATATATATAATTAAATA

TTTATAAATTATAATTTATAATATATTAAATATTTATATATATATATAT

ATATATATTAAACCATAGTTTTTTGATTCGTTTATAATATTTTAAA

ACTTTGTGAGTATATATATATATATATCTTATTCCCCCCCGGTGGTT

AATAAGTTTGGAGAGATATTATTTTATATTATGTATAAAA

**Fig.S2. Alignment of initiation context for A+T-rich region: there are some repeat in it, pane nucleotides indicate the repeat “T”, underlined nucleotides indicate the repeat “TA”.**
